# A Multiparametric and High-Throughput Platform for Host-Virus Binding Screens

**DOI:** 10.1101/2022.10.10.511545

**Authors:** Jan Schlegel, Bartlomiej Porebski, Luca Andronico, Leo Hanke, Steven Edwards, Hjalmar Brismar, Benjamin Murrell, Gerald McInerney, Oscar Fernandez-Capetillo, Erdinc Sezgin

## Abstract

Speed is key during infectious disease outbreaks. It is essential, for example, to identify critical host binding factors to the pathogens as fast as possible. The complexity of host plasma membrane is often a limiting factor hindering fast and accurate determination of host binding factors as well as high-throughput screening for neutralizing antimicrobial drug targets. Here we describe a multi-parametric and high-throughput platform tackling this bottleneck and enabling fast screens for host binding factors as well as new antiviral drug targets. The sensitivity and robustness of our platform was validated by blocking SARS-CoV-2 spike particles with nanobodies and IgGs from human serum samples.

**Teaser:** A fast screening platform tackling host-pathogen interactions.

## Introduction

Emerging microbial pathogens, such as bacteria, fungi and viruses, tremendously challenge human health and cause significant economical and societal burden worldwide. Therefore, tools facilitating and improving pandemic preparedness are of uttermost importance to minimize these negative effects. Current state-of-the-art methods, such as enzyme-linked immunosorbent assay (ELISA), reverse transcription-polymerase chain reaction (RT-PCR) and RT loop-mediated isothermal amplification (RT-LAMP) usually rely on bulk measurements resulting in a single readout-value (*1*). In addition, during the peaks of SARS-CoV-2 pandemic, RT-PCR instruments were used to capacity slowing down pandemic surveillance and highlighting the need for additional readout-systems. Especially flow cytometry, enabling fast and high-throughput measurements of complex mixtures, is widely used in clinics for immunophenotyping and would be an attractive and broadly available technique for such purposes (*2*).

To complement existing bulk measurement methods, we aimed to develop a fast and high-throughput platform to study host-pathogen interactions. The system should not only reconstitute host cell proteins, but also the lipid bilayer, which is mostly neglected in current state-of-the-art methods but often hosts important attachment factors. However, the complexity of the mammalian plasma membrane consisting of thousands of different lipids and proteins embedded between an outer glycocalyx and inner cortical cytoskeleton is overwhelming. This complexity not only slows down our efforts to identify important interaction partners but also obscures specific interactions between host and pathogen due to the plenitude of involved molecules and interactions. To overcome this bottleneck and reduce complexity, bottom-up model membrane systems are attractive alternatives which allow for precise control over composition and properties. Among these, planar supported lipid bilayer systems (SLBs) were widely used (*3*) but do not account for cells’ three-dimensional nature. Three-dimensional model systems, such as large unilamellar vesicles (LUVs), giant unilamellar vesicles (GUVs), and cell-derived giant plasma membrane vesicles (GPMVs) help to recreate cellular curvature but are challenging to use in high-throughput flow cytometry because of their fragility and size-inhomogeneity.

For this reason, we coated cell-sized 5μm silica beads with a lipid bilayer consisting of 98 mole percent 1-palmitoyl-2-oleoyl-glycero-3-phosphocholine (POPC) doped with 2 mole percent of a nickelated anchoring lipid (18:1 DGS-NTA(Ni)). Next, we attached His-tagged host-cell proteins of interest to membrane-coated beads to generate functionalized bead-supported lipid bilayers (fBSLBs) serving as minimal synthetic host-cells (Fig. 1A). In contrast to methods relying on random surface-adsorption, fBSLBs ensure proper protein orientation, tightly controllable receptor mobility and density as well as molecular interactions at the membrane plane. In addition, the presence of a hydrophobic lipid bilayer more closely mimics the cellular environment and enables to discriminate between binding preferences of pathogens to either host-cell proteins or lipids. For example, surface proteins of several viruses can bind different host-cell lipids facilitating cellular uptake and shaping viral tropism (*4*).

**Fig. 1.**
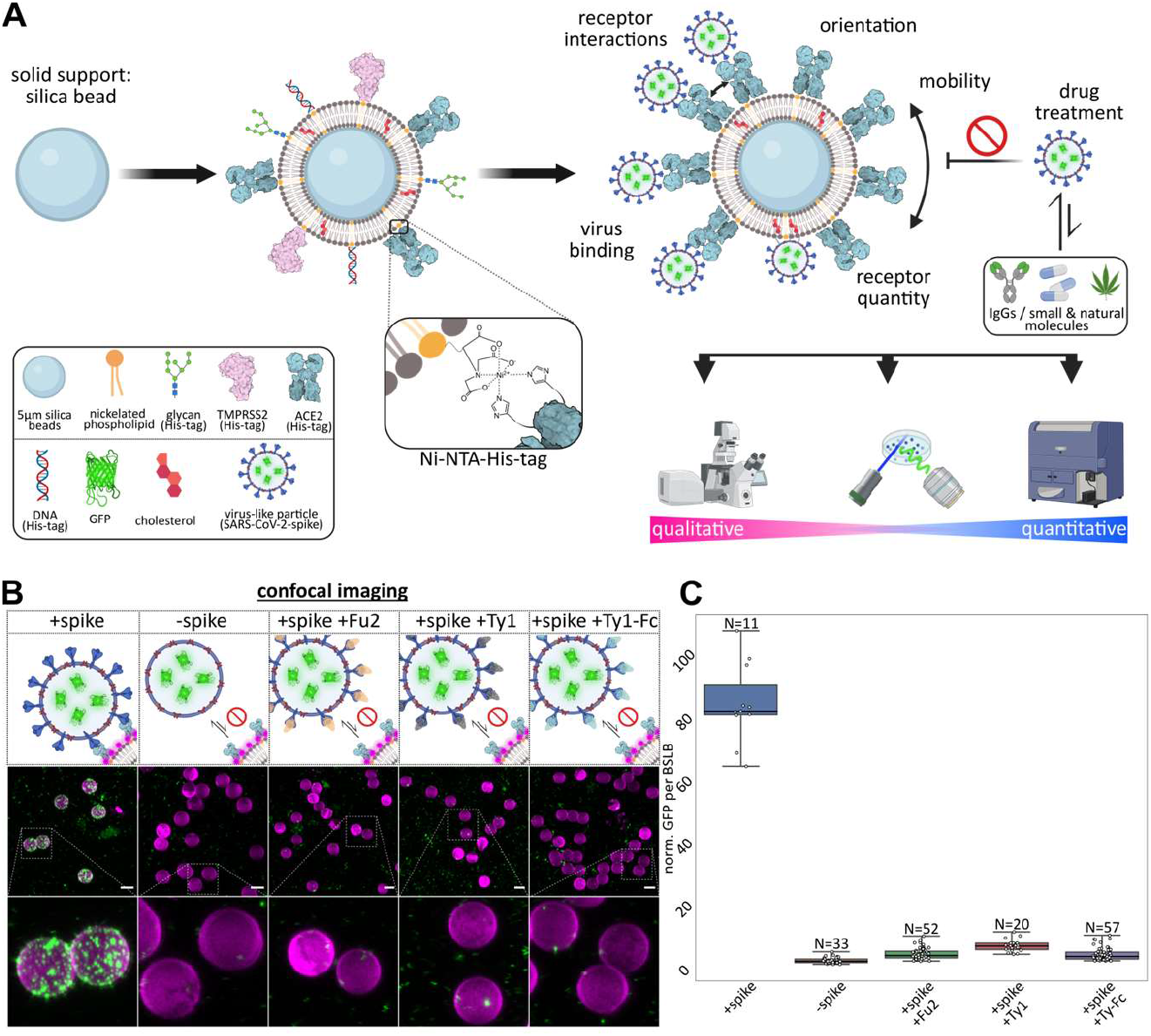
Design and characterization of our multi-parametric and high-throughput platform based on fBSLBs to study host-virus interactions. **(A)** Scheme depicting the bottom-up assembly of fBSLBs and available readout techniques. **(B)** LSM maximum-intensity projections of BSLBs (magenta) and VLPs (green) showing specific interaction between SARS-CoV-2 spike VLPs (+S-VLPs) and ACE2-fBSLBs. **(C)** Quantification of viral GFP-signal per fBSLBs of each condition from (b) shows specific attachment of +S-VLPs to ACE2-fBSLBs (median=84.05, N=11) and no interaction between -S-VLPs and ACE2-BSLBs (median=2.88, N=33) and nanobody-pretreated +S-VLPs and ACE2-BSLBs (Fu2: median=4.70, N=52; Ty1: median=7.77, N=20; Ty-Fc: median=4.41, N=57). Boxplot with overlay of individual data points, median as black center line, box showing the quartiles and whiskers from minimum to maximum value. Illustrations were created using Biorender.com and Inkscape.

In this study, we show that fBSLBs carrying different host cell receptors, such as angiotensin-converting enzyme 2 (ACE2), can serve as highly diverse platform to screen for unknown protein- and lipid-binding molecules, drugs influencing host-pathogen interactions and the blocking efficiency of neutralizing antibodies present in human serum samples. Its fast implementation, easy adaptability of multiple parameters and high-throughput capability propel our method as an important platform to understand and tackle host-pathogen interactions.

## Results

### fBSLBs enable fast and qualitative host-pathogen interaction studies

Upon coating of 5μm silica beads with POPC:DGS-NTA(Ni) 98:2 mole percent of liposome solution, we verified proper bilayer formation by measuring diffusion of a fluorescent lipid analogue using fluorescence correlation spectroscopy (FCS) (fig. S1, A and B), which matched with previous data (*5*, *6*). We first generated fBSLBs carrying ACE2 and studied their interaction with SARS-CoV-2 spike expressing virus-like particles (+S-VLPs) using confocal microscopy (Fig.1, B and C). To quantify VLP-binding per bead we developed an automated image analysis workflow using Fiji (*7*) (fig. S2). While there was strong interaction between ACE2-fBSLBs and +S-VLPs, it was absent in VLPs with no spike (-S-VLPs) and +S-VLPs pre-treated with SARS-CoV-2 neutralizing spike nanobodies which were shown to be potent tools to neutralize SARS-CoV-2 by blocking the interaction between spike receptor-binding domain (RBD) and its host receptor ACE2 (*8*, *9*). Thus, fBSLBs can serve as powerful screening platform to identify efficient inhibitors with therapeutic potential.

### fBSLBs enable quantitative high-throughput screening

To increase number of data points and decrease acquisition time, we performed fast, quantitative, 3D lattice light-sheet microscopy (LLSM) and quantified viral loads per fBSLB (Fig. 2, A and B) which confirmed confocal microscopy data. To screen several tens of thousands of fBSLBs within minutes, fast and high-throughput flow cytometry can be used thanks to the firm nature of fBSLBs. Individual fBSLBs were easily detected by their specific scattering signal and presence of the lipid bilayer confirmed by 1,2-dioleoyl-sn-glycero-3-phosphoethanolamine Abberior STAR RED (ASR-PE) labelling while VLPs were labelled with eGFP. Upon addition of ASR-PE and +S-VLPs to ACE2-fBSLBs, we observed a strong increase of fluorescence intensity per bead both in virus (green) and in membrane (red) channels (Fig. 2C). Moreover, virus signal decreased significantly upon nanobody treatment, confirming the neutralizing ability of nanobodies. Hence, fBSLBs enable to study host-virus interactions using quantitative high-throughput flow cytometry which is usually not feasible due to the small size of viral particles. Moreover, it serves as a powerful platform to study concentration-dependent effects of molecules on the binding between viruses and host-cell receptors. To show this, we determined optimal concentrations of ACE2 on the fBSLBs and the amount +S-VLPs by titration series (fig. S3).

**Fig. 2.**
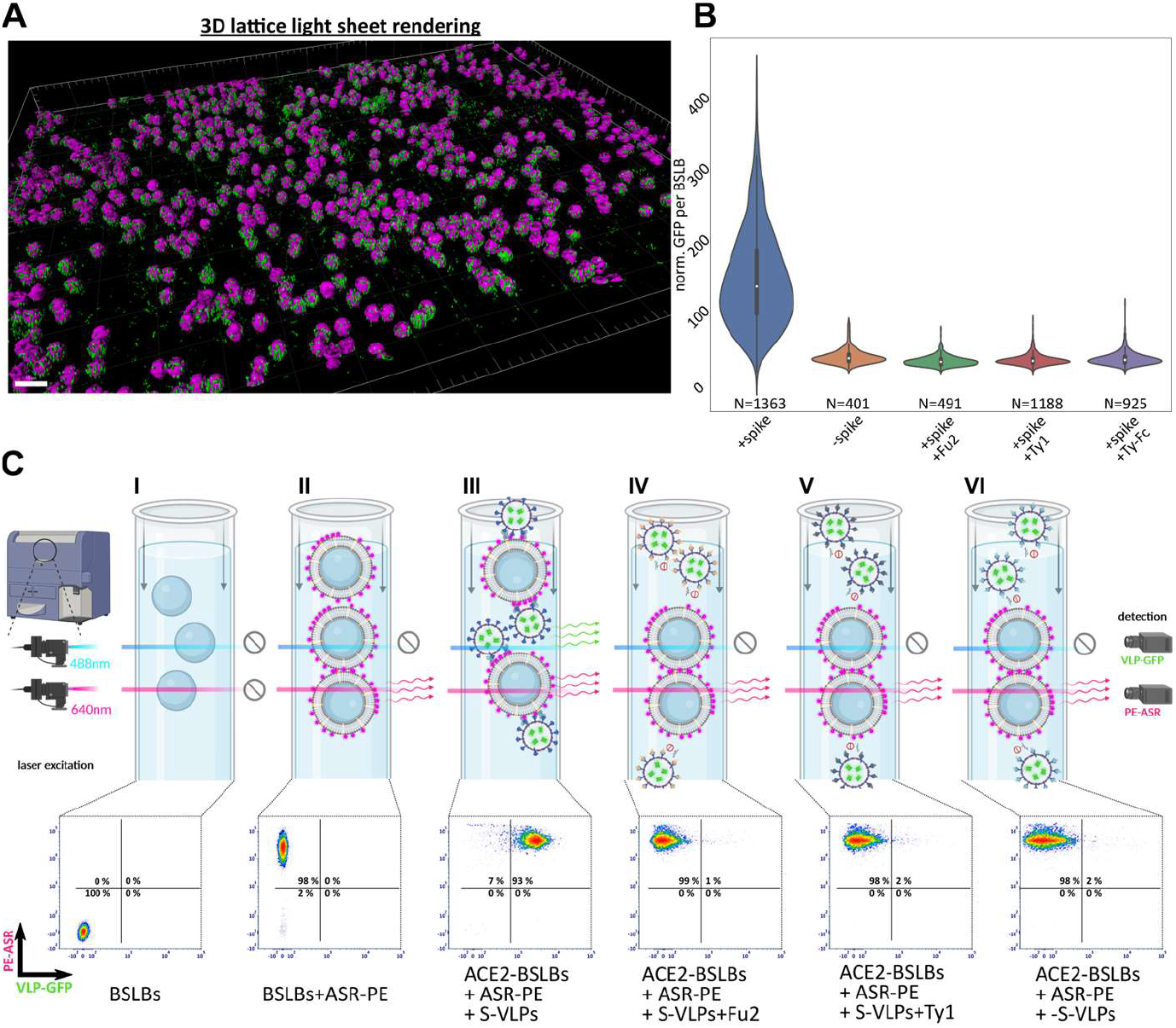
High-throughput measurements using fBSLBs. **(A)** Fast and quantitative LLSM enabling big-volume renderings of ACE2-fBSLBs (magenta) interacting with +S-VLPs (green). **(B)** Quantification of VLP-GFP signal per bead proves specific interaction between +S-VLPs and ACE2-fBSLBs (N>400). Violin plot with miniature boxplot showing quartiles and median as white dot. +S-VLPs show significant increased binding to ACE2-fBSLBs as compared to the other groups (p-value<0.0001). **(C)** Fast high-throughput screening of interaction between VLPs and ACE2-fBSLBs using flow cytometry. Strong signal of the fluorescent lipid ASR-PE (y-axis) confirms functional bilayer formation and interaction of VLPs with fBSLBs can be followed by intensity changes in the VLP-GFP channel (x-axis) (N>8500 per condition). Illustrations were created using Biorender.com and Inkscape.

### Screening receptors using fBSLBs

Besides ACE2, other receptors have been described to contribute to SARS-CoV-2 binding to the host-cell surface and subsequent infection. For this reason, we tested interaction of +S-VLPs with reported host-cell receptors Neuropilin-1 (CD304) (*10, 11*), Basigin (CD147) (*12*), DPP4 (CD26) (*13, 14*) and TMPRSS2 (*15*, *16*) using fBSLBs in combination with flow cytometry. As expected, +S-VLPs showed strongest interaction with ACE2-fBSLBs (Fig. 3A). Interestingly, +S-VLPs also interacted with CD304-fBSLBs and TMPRSS2-fBSLBs, confirming that these two proteins act as host binding factors, but neither interaction was as strong as for ACE2-fBSLBs. No binding was observed for CD147-fBSLBs or CD26-fBSLBs, suggesting that these proteins cannot act as host binding factors alone and might require additional host-cell binding elements. Notably, CD304-fBSLBs binding to VLPs was independent of spike-protein on their surface, e.g., -S-VLPs also bound to CD304-fBSLBs effectively while they did not bind any other proteins we tested (Fig. 3B). This suggests the presence of another interaction partner on the viral particles to this receptor. To check this hypothesis, we performed dual-receptor screens with each individual receptor in absence or presence of same molar concentration of ACE2 (Fig. 3C). The presence of ACE2 always significantly increased the interaction of +S-VLPs with fBSLBs, but the overall strongest binding was observed in the simultaneous presence of CD304 and ACE2, supporting the idea of two different additive binding mechanisms.

**Fig. 3.**
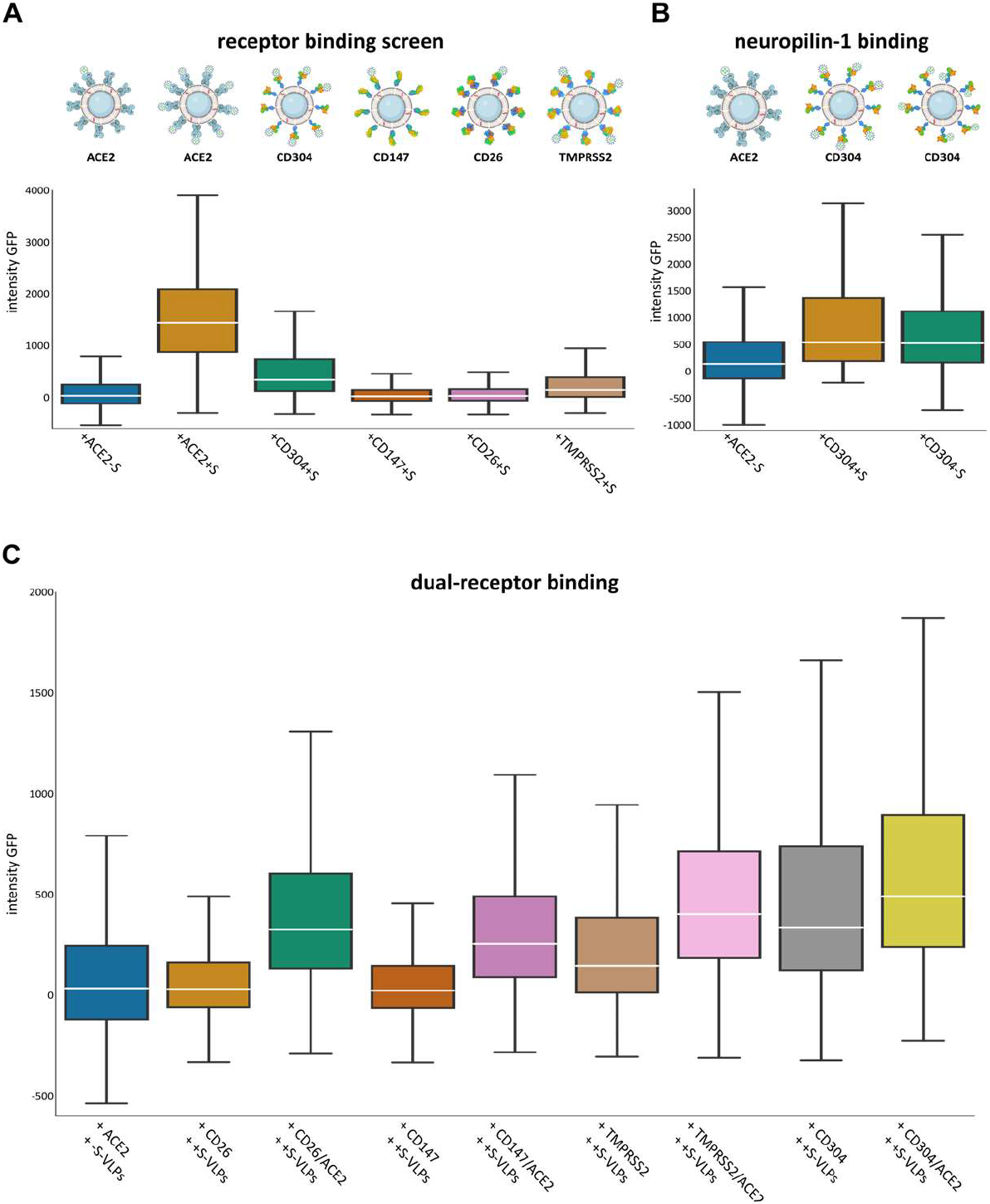
Receptor screening using fBSLBs. **(A)** Application of fBSLBs to study interaction with different host-receptors. Scheme depicting fBSLBs with different His-tagged host-receptors. Box-and-whisker plots showing the distribution of +S-VLP-GFP signal of 40000 BSLBs analyzed by flow cytometry. Besides ACE2 (median: 1440), specific but less pronounced binding was also observed for CD304 (median: 337) and TMPRSS2 (median: 145). Except for the dataset-pair +ACE2-S (blue) and +CD26+S (magenta) all populations are significantly different from each other (p-value<0.0001). **(B)** Interaction of VLPs with CD304-fBSLBs in the absence of spike protein (N=20000). All populations are significantly different from each other (p-value<0.0001). **(C)** Dual receptor screen using fBSLBs and flow cytometry. No interaction between +S-VLPs and the host-cell receptors CD26 (median: 31) and CD147 (median: 23) was detected, respectively. Increased interaction with TMPRSS2 (median: 145) and CD304 (Neuropilin-1, median: 337) fBSLBs was observed, respectively. Upon coating BSLBs with 1:1 molar ratio of ACE2 and different host-cell receptors all interactions were further increased while the receptor pairs TMPRSS2/ACE2 (median: 403) and CD304/ACE2 (median: 491) showed strongest binding of +S-VLPs (N=40000 per condition). All populations are significantly different from each other (p-value<0.0001). Box plots show inter-quartile range with white median line and whiskers extend to 1.5 inter-quartile range.

fBSLBs allow tight control not only on the composition of surface proteins but also of lipid composition. We made use of this and screened for reported lipid co-receptors for spike, such as GM1 gangliosides (*17*). Despite varying GM1 concentrations in fBSLBs, we could not observe any concentration-dependent binding of VLPs pseudotyped with spike, beta-spike, delta-spike, Ebola virus glycoprotein (GP) or without any viral protein (fig. S4, A and B). These results highlight the need for additional high-affinity host-cell binding factors for efficient virus-host interaction.

### Surveillance of human serum samples using fBSLBs

Key for pandemic containment is surveillance of convalescent serum samples and their ability to block the interaction between virus and host cell receptors. Virus-specific antibody levels in human serum are usually proportional to neutralization of the virus and can be used to predict disease-outcome or the need for additional booster vaccinations (*18, 19*). Moreover, it is very important to understand whether anti-viral IgGs in prevalent serum samples still protect from upcoming new variants to decide for vaccine-adjustments and therapeutic treatment options. To show the potential of our method to answer these questions, we first determined the amount of spike-IgGs in three human serum samples using a bead-based assay in combination with flow cytometry (**Fig. 4, upper panel**). Glass beads were coated with recombinant spike receptor binding domain (RBD), incubated with serum samples, and anti-spike IgGs detected by labelling with secondary dye-conjugated anti-human antibodies. After we determined the relative levels of anti-spike IgGs in the three serum samples, we blocked +S-VLPs with the different serum samples and studied the interaction with ACE2-fBSLBs. The amount of anti-spike IgGs perfectly correlated with the blocking efficiency, highlighting the ability of this method as powerful tool for pandemic surveillance (**Fig. 4, lower panel**).

**Fig. 4.**
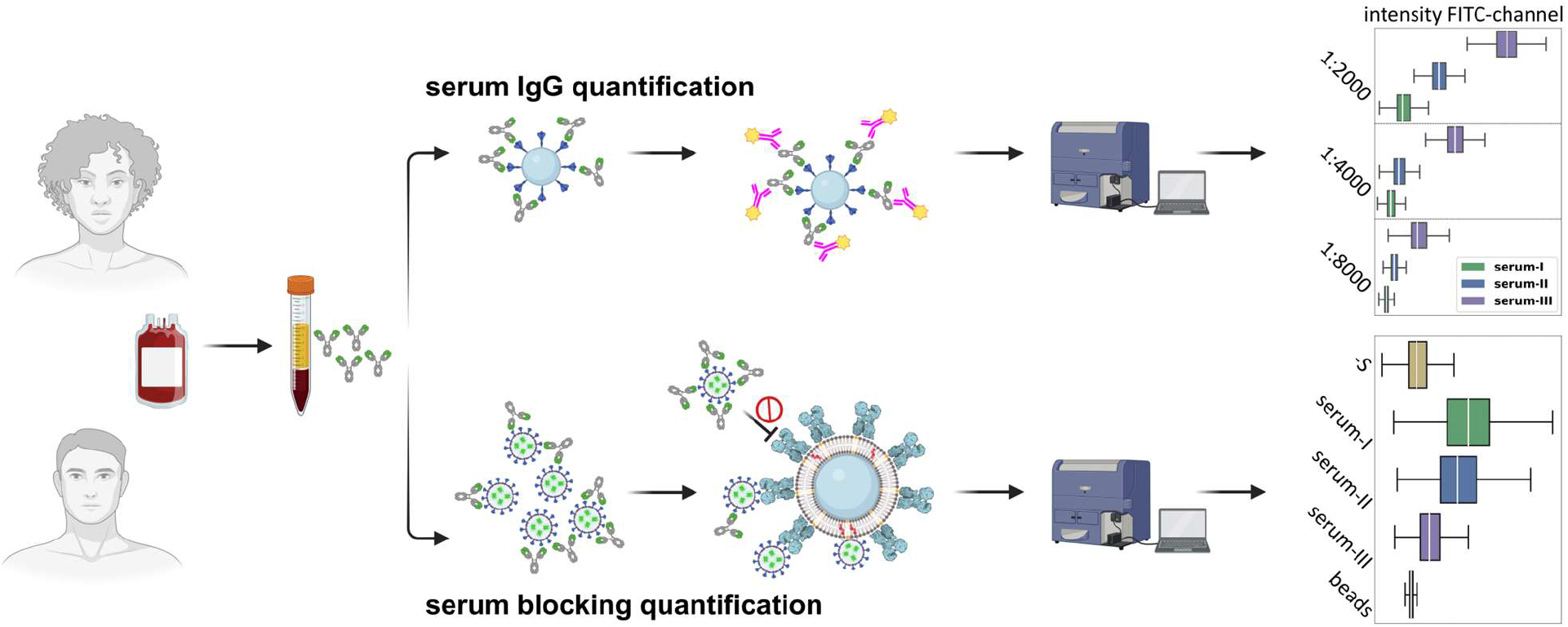
Application of fBSLBs to study blocking efficacy of neutralizing antibodies in human blood serum samples. Scheme showing the processing of human serum samples to quantify amount of anti-spike IgGs and their capacity to block the interaction between +S-VLPs and ACE2-fBSLBs. Upper box plots illustrating the amount of anti-spike IgGs in three different serum samples at different dilutions (N=10000). Lower box plots showing the efficiency to block the interaction between +S-VLPs and ACE2-BSLBs for the three different serum samples of each population (N=10000). Note that the amount of anti-spike-IgGs in the serum samples correlates with blocking efficiency. All populations are significantly different from each other (p-value<0.0001). Box plots show inter-quartile range with white median line and whiskers extend to 1.5 inter-quartile range. Illustrations were created using Biorender.com and Inkscape.

## Discussion

Quick response to pandemic outbreaks is of uttermost importance for disease and damage control. Our platform relies on material and molecules which are available from early pandemic onset, such as the sequence of viral structural proteins and potential interaction partners. Exploiting highly specific Ni-NTA-His-tag conjugation makes the platform highly versatile and accessible, since this chemistry is widely used for protein purification and His-tagged proteins are available from a myriad of commercial resources. Screening of potential host-cell receptors and co-receptors, including lipids, can be done within a few hours using qualitative and high-throughput quantitative readout platforms. In contrast to other methods, our method enables tight control of multiple cellular parameters such as lipid composition, receptor mobility, receptor orientation, receptor-receptor interactions, and local receptor densities. The platform allows to determine serum-virus neutralization capacity in a safe laboratory environment within hours. Moreover, the presence of a lipid bilayer more closely mimics the cellular environment and can help to entangle the complex interplay between virus-receptor and virus-bilayer interactions which are often difficult to discriminate. Due to its highly defined bottom-up assembly, fBSLBs are not prone to cellular heterogeneity, e.g. due to differences in cell-cycle states, transcription and translation, which can complicate drug screens.

However, this cellular heterogeneity could fine-tune host-pathogen interactions which is challenging to reproduce with our platform. Recent advances on coating beads with native cellular membranes would be an opportunity to recreate this complexity (*20, 21*). Another limitation of fBSLBs is its inability to initially detect cellular toxic compounds. This can also be advantageous since substances showing both cellular toxicity and binding inhibition are identified and not directly discarded. Further efforts in reducing cellular toxicity while maintaining inhibitory effects of these molecules would be an exciting way to find new drug targets.

Our broadly accessible platform enables to perform fast and high-throughput drug screens and to discriminate whether drugs act on the virus particles or on the host-cell receptors. Due to its bottom-up design, our method should be readily extensible to other biomolecules (e.g. glycocalyx, DNA, RNA) and pathogens (bacteria, fungi) making it a valuable tool for future pandemic preparedness.

## Materials and Methods

### fBSLB preparation

fBSLBs were prepared similar as described previously (*6*). For preparation of one batch of fBSLBs, 1×10^7^ 5μm silica beads (Bangs Laboratories) were vortexed thoroughly and washed three times with PBS using 1500xg and 30 seconds centrifugation steps. Beads were coated with lipid bilayers of defined compositions by incubation with 100μl 0.5mg/ml liposomes shaking at 1400rpm for 30 minutes. Liposomes were formed by mixing lipids dissolved in chloroform, solvent evaporation under a steam of nitrogen, re-hydration, and tip-sonication (Branson Sonifier 250). To prepare fBSLBs with His-tagged proteins, a lipid mixture consisting of 98mol% 16:0-18:1 POPC and 2mol% 18:1 DGS-Ni:NTA (Avanti Polar Lipids) was used. After bilayer formation beads were washed two more times with PBS and 5pmol of His-tagged proteins added (Sino Biological: ACE2-His 10108-H08H, Neuropilin-1-His 10011-H08H, CD147-His 10186-H08H, CD26-His 10688-H08H). After 20 minutes on a rotary shaker the bilayer of fBSLBs was optionally directly labelled with a fluorescent lipid analogue followed by 2 washing steps with PBS. Final fBSLBs were diluted in 500μl PBS and used the same day. To study host-virus interactions, 20μl of fBSLBs were mixed with 15μl of GFP-tagged pseudotyped VLPs and incubated for 30 minutes on a rotary shaker at room temperature and directly used for microscopy or flow cytometry. Optionally, VLPs were pre-treated for 20 minutes on ice with 2μM Ty1, Ty1-Fc or Fu2 nanobodies4,5.

### VLP preparation

Mycoplasma-free HEK293T cells were cultured in DMEM supplemented with 10% FCS and grown to ~70% confluency in T75 cell culture flasks. To produce VLPs, cells were co-transfected using Lipofectamine 3000 and 15μg of DNA encoding for viral protein (pCMV14-3X-Flag-SARS-CoV-2 S was a gift from Zhaohui Qian - Addgene plasmid # 145780; delta/beta spike expression plasmid kindly provided by Benjamin Murrell; Ebola GP expression plasmid kindly provided by Jochen Bodem), 7.5μg DNA encoding for HIV Vpr-GFP (NIH HIV Reagent Program, Division of AIDS, NIAID, NIH: pEGFP-Vpr, ARP-11386, contributed by Dr. Warner C. Greene), and 7.5μg encoding for a lentiviral packaging plasmid (psPAX2 was a gift from Didier Trono - Addgene plasmid # 12260). Media was exchanged after 12 hours and VLPs harvested after 24 and 48 hours and enriched fiftyfold using LentiX concentrator according to the protocol provided by the manufacturer (Takara).

### Microscopy and Quantification

After incubation with pseudotyped VLPs, fBSLBs were put into chambered glass coverslips (IBIDI: 81817) and imaging performed in PBS. Confocal microscopy was performed using a C-Apochromat 40x/1.20 water immersion objective of the Zeiss LSM780 microscope. Viral GFP was excited using 488nm argon laser and membrane-inserted ASR-PE was excited using a 633nm helium neon laser, while emission was collected from 498-552nm and 641-695nm, respectively. Full surface of 5μm fBSLBs was recorded by acquiring z-stacks with 24 slices each 0.3μm and VLP-GFP signal per bead quantified using ImageJ following the provided macro and automated workflow of Suppl. Fig. 02. To acquire fast, gentle, and big 3D volumes we used LLSM (Zeiss Lattice Lightsheet 7) with 488nm and 640nm laser excitation for viral GFP and ASR-PE, respectively. The general analysis workflow followed the one for confocal data, but parameters were adjusted for differences in signal intensity.

### Flow Cytometry

Upon interaction of VLPs with fBSLBs the mixture was diluted in 500μl PBS and transferred into flow tubes. Flow cytometry was performed using a BD Fortessa system acquired at low speed and 488nm (FITC) or 640nm (APC) excitation/emission settings used for VLP-GFP and ASR-PE, respectively. 10 000 to 20 000 events were acquired and analysed using FCS Express 7 and Python (FCSParser). Gating was only performed for data shown in Fig. 1f on singlet bead population clearly visible in the forward-versus side-scatter plot. This population was always at least 85% of the total bead population.

### Serum Blocking

Human blood from healthy donors was obtained from blood transfusion station of Karolinska Hospital and serum prepared by centrifugation. The serum was aliquoted and frozen for further later use. To determine the amount of anti-spike IgGs in serum samples, 1×107 5μm silica beads (Bangs Laboratories) were washed three times with PBS and coated for 30 minutes with 47pmol SARS-CoV-2 RBD (BioSite: 40592-V08H) on a rotary shaker. After two washing steps with PBS beads were resuspended in 500μl PBS supplemented with 4mg/ml BSA to block non-specific interaction sites. 20μl of beads were incubated with stated serum dilutions over night at 4°C on a rotary shaker to enable interaction of anti-spike IgGs with coated beads. After two washing steps, anti-spike IgGs were labelled by incubation with 4μg/ml secondary anti-human IgG Alexa Fluor 488 antibodies (ThermoFischer: A11013) for one hour at room temperature on a rotary shaker in the dark. Labelled beads were washed and signal intensity of at least 9000 beads determined by flow cytometry. To test serum blocking efficiency, VLPs were pre-treated with stated serum concentrations over night at 4°C on a rotary shaker before incubated with ACE2-fBSLBs as described above.

### Statistical Analysis

Visualization and statistical analysis of the data was performed using Python (Anaconda Navigator 2.3.2, JupyterLab 3.2.9) and Kruskal-Wallis H-test with post hoc pairwise test for multiple comparisons (Dunn’s test with Bonferroni one-step correction). Standard error of the median was estimated by multiplying the standard error of the mean with the constant 1.253.

## Acknowledgments

We appreciate the contribution of the National Microscopy Infrastructure, SciLifeLab COVID-19 Research Program, European Union’s Horizon 2020 research and innovation program, G2P-UK National Virology consortium and Barclay Lab at Imperial College for providing the plasmids Beta/B.1.351 and Delta/B.1.617.2. We thank Jaromir Mikes for support with flow cytometry

## Funding

National Microscopy Infrastructure, NMI, VR-RFI 2016-00968

Knut and Alice Wallenberg Foundation and SciLifeLab, COVID-19 Research Program

European Union’s Horizon 2020 research and innovation program, 101003653 (CoroNAb)

G2P-UK National Virology consortium, MRC/UKRI, MR/W005611/1

## Author contributions

Conceptualization: ES, JS

Methodology: ES, JS, BP, LA, LH, SE

Investigation: ES, JS,

Visualization: ES, JS

Supervision: ES, HB, BM, GM, OFC

Writing—original draft: JS, ES

Writing—review & editing: JS, ES, BP, LA, LH, BM, GM, OFC

## Competing interests

Authors declare that they have no competing interests

## Data and materials availability

All raw data will be available upon publication (FigShare DOI: 10.17044/scilifelab.20517336).

## Notes

### Competing Interest Statement

The authors have declared no competing interest.

### Summary of Updates

The manuscript has been re-structured. Instead of 2 figures, we have now 4 figures.

## References

1. B. D. Kevadiya, J. Machhi, J. Herskovitz, M. D. Oleynikov, W. R. Blomberg, N. Bajwa, D. Soni, S. Das, M. Hasan, M. Patel, A. M. Senan, S. Gorantla, J. McMillan, B. Edagwa, R. Eisenberg, C. B. Gurumurthy, S. P. M. Reid, C. Punyadeera, L. Chang, H. E. Gendelman, Diagnostics for SARS-CoV-2 infections. Nat. Mater. 20, 593–605 (2021).

2. H. T. Maecker, J. P. McCoy, R. Nussenblatt, Standardizing immunophenotyping for the Human Immunology Project. Nat Rev Immunol. 12, 191–200 (2012).

3. T. Sych, C. O. Gurdap, L. Wedemann, E. Sezgin, How Does Liquid-Liquid Phase Separation in Model Membranes Reflect Cell Membrane Heterogeneity? Membranes. 11, 323 (2021).

4. M. Mazzon, J. Mercer, Lipid interactions during virus entry and infection: Lipids and Viruses. Cell Microbiol. 16, 1493–1502 (2014).

5. D. Beckers, D. Urbancic, E. Sezgin, Impact of Nanoscale Hindrances on the Relationship between Lipid Packing and Diffusion in Model Membranes. J. Phys. Chem. B. 124, 1487–1494 (2020).

6. P. F. Céspedes, A. Jainarayanan, L. Fernández-Messina, S. Valvo, D. G. Saliba, E. Kurz, A. Kvalvaag, L. Chen, C. Ganskow, H. Colin-York, M. Fritzsche, Y. Peng, T. Dong, E. Johnson, J. A. Siller-Farfán, O. Dushek, E. Sezgin, B. Peacock, A. Law, D. Aubert, S. Engledow, M. Attar, S. Hester, R. Fischer, F. Sánchez-Madrid, M. L. Dustin, T-cell trans-synaptic vesicles are distinct and carry greater effector content than constitutive extracellular vesicles. Nat Commun. 13, 3460 (2022).

7. J. Schindelin, I. Arganda-Carreras, E. Frise, V. Kaynig, M. Longair, T. Pietzsch, S. Preibisch, C. Rueden, S. Saalfeld, B. Schmid, J.-Y. Tinevez, D. J. White, V. Hartenstein, K. Eliceiri, P. Tomancak, A. Cardona, Fiji: an open-source platform for biological-image analysis. Nat Methods. 9, 676–682 (2012).

8. L. Hanke, L. Vidakovics Perez, D. J. Sheward, H. Das, T. Schulte, A. Moliner-Morro, M. Corcoran, A. Achour, G. B. Karlsson Hedestam, B. M. Hällberg, B. Murrell, G. M. McInerney, An alpaca nanobody neutralizes SARS-CoV-2 by blocking receptor interaction. Nat Commun. 11, 4420 (2020).

9. L. Hanke, H. Das, D. J. Sheward, L. Perez Vidakovics, E. Urgard, A. Moliner-Morro, C. Kim, V. Karl, A. Pankow, N. L. Smith, B. Porebski, O. Fernandez-Capetillo, E. Sezgin, G. K. Pedersen, J. M. Coquet, B. M. Hällberg, B. Murrell, G. M. McInerney, A bispecific monomeric nanobody induces spike trimer dimers and neutralizes SARS-CoV-2 in vivo. Nat Commun. 13, 155 (2022).

10. L. Cantuti-Castelvetri, R. Ojha, L. D. Pedro, M. Djannatian, J. Franz, S. Kuivanen, F. van der Meer, K. Kallio, T. Kaya, M. Anastasina, T. Smura, L. Levanov, L. Szirovicza, A. Tobi, H. Kallio-Kokko, P. Österlund, M. Joensuu, F. A. Meunier, S. J. Butcher, M. S. Winkler, B. Mollenhauer, A. Helenius, O. Gokce, T. Teesalu, J. Hepojoki, O. Vapalahti, C. Stadelmann, G. Balistreri, M. Simons, Neuropilin-1 facilitates SARS-CoV-2 cell entry and infectivity. Science. 370, 856–860 (2020).

11. J. L. Daly, B. Simonetti, K. Klein, K.-E. Chen, M. K. Williamson, C. Antón-Plágaro, D. K. Shoemark, L. Simón-Gracia, M. Bauer, R. Hollandi, U. F. Greber, P. Horvath, R. B. Sessions, A. Helenius, J. A. Hiscox, T. Teesalu, D. A. Matthews, A. D. Davidson, B. M. Collins, P. J. Cullen, Y. Yamauchi, Neuropilin-1 is a host factor for SARS-CoV-2 infection. Science. 370, 861–865 (2020).

12. K. Wang, W. Chen, Z. Zhang, Y. Deng, J.-Q. Lian, P. Du, D. Wei, Y. Zhang, X.-X. Sun, L. Gong, X. Yang, L. He, L. Zhang, Z. Yang, J.-J. Geng, R. Chen, H. Zhang, B. Wang, Y.-M. Zhu, G. Nan, J.-L. Jiang, L. Li, J. Wu, P. Lin, W. Huang, L. Xie, Z.-H. Zheng, K. Zhang, J.-L. Miao, H.-Y. Cui, M. Huang, J. Zhang, L. Fu, X.-M. Yang, Z. Zhao, S. Sun, H. Gu, Z. Wang, C.-F. Wang, Y. Lu, Y.-Y. Liu, Q.-Y. Wang, H. Bian, P. Zhu, Z.-N. Chen, CD147-spike protein is a novel route for SARS-CoV-2 infection to host cells. Sig Transduct Target Ther. 5, 283 (2020).

13. N. Vankadari, J. A. Wilce, Emerging COVID-19 coronavirus: glycan shield and structure prediction of spike glycoprotein and its interaction with human CD26. Emerging Microbes & Infections. 9, 601–604 (2020).

14. Y. Li, Z. Zhang, L. Yang, X. Lian, Y. Xie, S. Li, S. Xin, P. Cao, J. Lu, The MERS-CoV Receptor DPP4 as a Candidate Binding Target of the SARS-CoV-2 Spike. iScience. 23, 101160 (2020).

15. M. Hoffmann, H. Kleine-Weber, S. Schroeder, N. Krüger, T. Herrler, S. Erichsen, T. S. Schiergens, G. Herrler, N.-H. Wu, A. Nitsche, M. A. Müller, C. Drosten, S. Pöhlmann, SARS-CoV-2 Cell Entry Depends on ACE2 and TMPRSS2 and Is Blocked by a Clinically Proven Protease Inhibitor. Cell. 181, 271–280.e8 (2020).

16. M. Hussain, N. Jabeen, A. Amanullah, A. Ashraf Baig, B. Aziz, S. Shabbir, F. Raza, N. Uddin, 1 Bioinformatics and Molecular Medicine Research Group, Dow Research Institute of Biotechnology and Biomedical Sciences, Dow College of Biotechnology, Dow University of Health Sciences, Karachi-Pakistan, 2 Department of Microbiology, University of Karachi, Karachi-Pakistan, 3 Faculty of Computer Science, IBA, Karachi-Pakistan, Molecular docking between human TMPRSS2 and SARS-CoV-2 spike protein: conformation and intermolecular interactions. AIMS Microbiology. 6, 350–360 (2020).

17. L. Nguyen, K. A. McCord, D. T. Bui, K. M. Bouwman, E. N. Kitova, M. Elaish, D. Kumawat, G. C. Daskhan, I. Tomris, L. Han, P. Chopra, T.-J. Yang, S. D. Willows, A. L. Mason, L. K. Mahal, T. L. Lowary, L. J. West, S.-T. D. Hsu, T. Hobman, S. M. Tompkins, G.-J. Boons, R. P. de Vries, M. S. Macauley, J. S. Klassen, Sialic acid-containing glycolipids mediate binding and viral entry of SARS-CoV-2. Nat Chem Biol. 18, 81–90 (2022).

18. D. S. Khoury, D. Cromer, A. Reynaldi, T. E. Schlub, A. K. Wheatley, J. A. Juno, K. Subbarao, S. J. Kent, J. A. Triccas, M. P. Davenport, Neutralizing antibody levels are highly predictive of immune protection from symptomatic SARS-CoV-2 infection. Nat Med. 27, 1205–1211 (2021).

19. T. A. Bates, H. C. Leier, Z. L. Lyski, S. K. McBride, F. J. Coulter, J. B. Weinstein, J. R. Goodman, Z. Lu, S. A. R. Siegel, P. Sullivan, M. Strnad, A. E. Brunton, D. X. Lee, A. C. Adey, B. N. Bimber, B. J. O’Roak, M. E. Curlin, W. B. Messer, F. G. Tafesse, Neutralization of SARS-CoV-2 variants by convalescent and BNT162b2 vaccinated serum. Nat Commun. 12, 5135 (2021).

20. S. K. Cheppali, R. Dharan, R. Katzenelson, R. Sorkin, ACS Appl. Mater. Interfaces, in press, doi:10.1021/acsami.2c13095.

21. L. Liu, D. Pan, S. Chen, M.-V. Martikainen, A. Kårlund, J. Ke, H. Pulkkinen, H. Ruhanen, M. Roponen, R. Käkelä, W. Xu, J. Wang, V.-P. Lehto, Systematic design of cell membrane coating to improve tumor targeting of nanoparticles. Nat Commun. 13, 6181 (2022).

