## Supplementary Figures for "A Multiparametric and High-Throughput Platform for Host-Virus Binding Screens"

Jan Schlegel *et al.*

*Corresponding author: Erdinc Sezgin


**This PDF file includes:**

Figs. S1 to S4

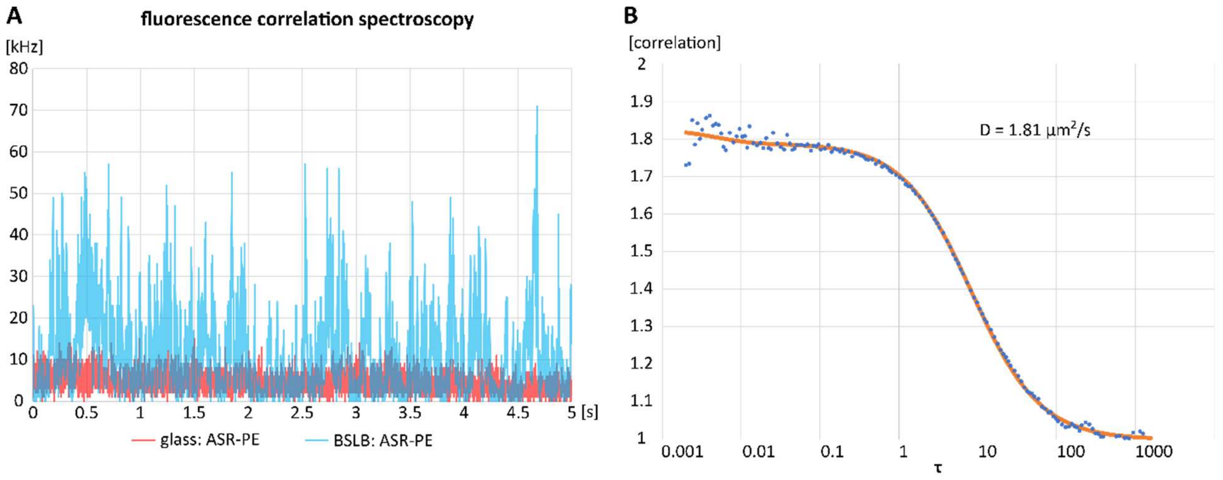


**Fig. S1.**

**Proof of functional lipid-bilayer formation. A.** Raw trace of FCS measurement of the lipid bilayer marker (ASR-PE) attached to 5μm silica beads in the presence (cyan) or absence (red) of the used lipid bilayer (98mol% POPC & 2mol% 18:1 DGS-Ni:NTA). Attachment and mobility of ASR-PE to the silica beads was only observed in the presence of a lipid bilayer (cyan), as derived from the fluorescence signal (y-axis: count rate) and fluctuations (x-axis: measurement time), respectively. **B.** Diffusion coefficient of 1.81μm$^2$/s for ASR-PE in BSLB was obtained by fitting (orange curve) of FCS-data (blue dots) with FoCuS-point software.

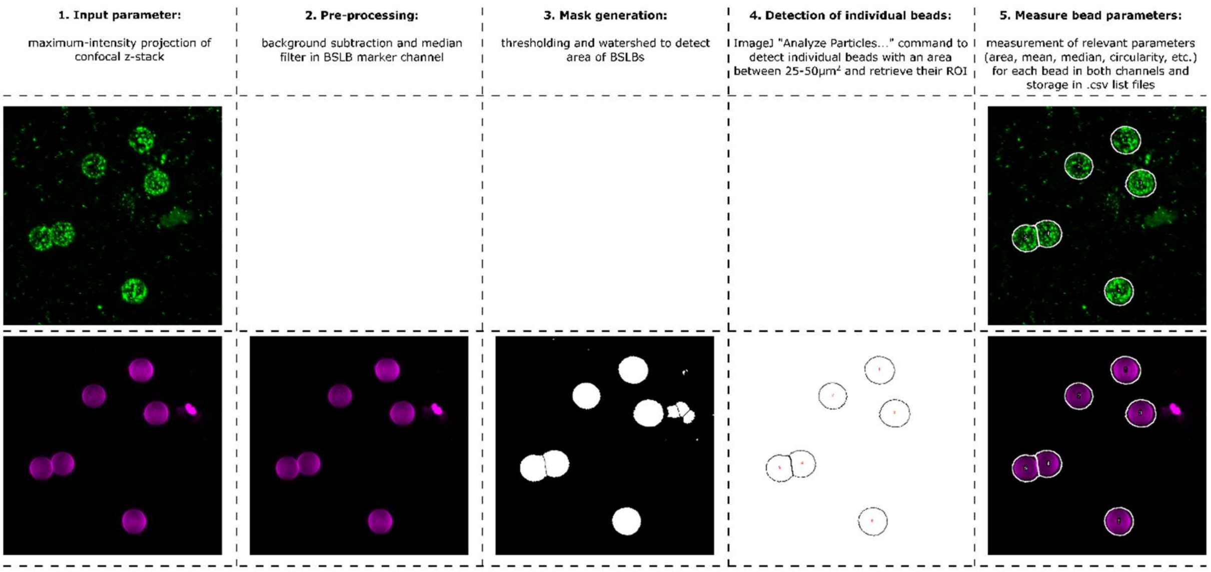


**Fig. S2.**

**Automated image analysis workflow to determine VLP-GFP signal per fBSLB in confocal and LLSM data.** Maximum-intensity projections of z-stacks were processed and thresholded to retrieve the region of interests (ROIs) of individual fBSLBs in the lipid-marker channel. ROIs were analyzed in the viral GFP-channel to extract mean and median intensity values per fBSLB.

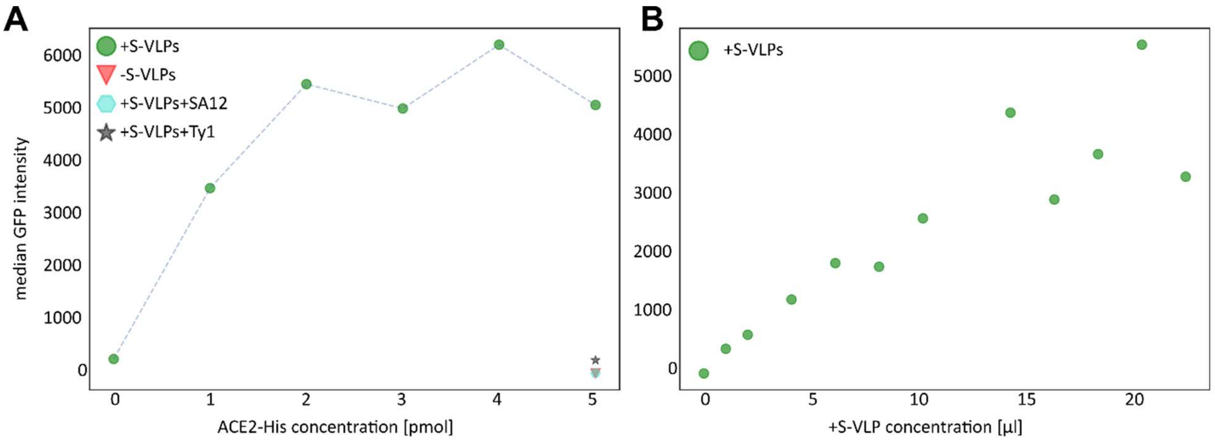


**Fig. S3.**

**Optimal protein- and VLP-concentrations for fBSLB were determined by titration experiments combined with flow cytometry. A.** 10x10$^6$ BSLBs were functionalized with increasing amounts of ACE2-His protein and incubated with constant amount of +S-VLPs. Each dot in the scatterplot shows the median GFP-intensity of >20000 fBSLBs. **B.** 10x10$^6$ BSLBs were functionalized with constant amount of ACE2-His protein and incubated with increasing concentrations of +S-VLPs. Each dot in the scatterplot shows the median GFP-intensity of >20000 fBSLBs.

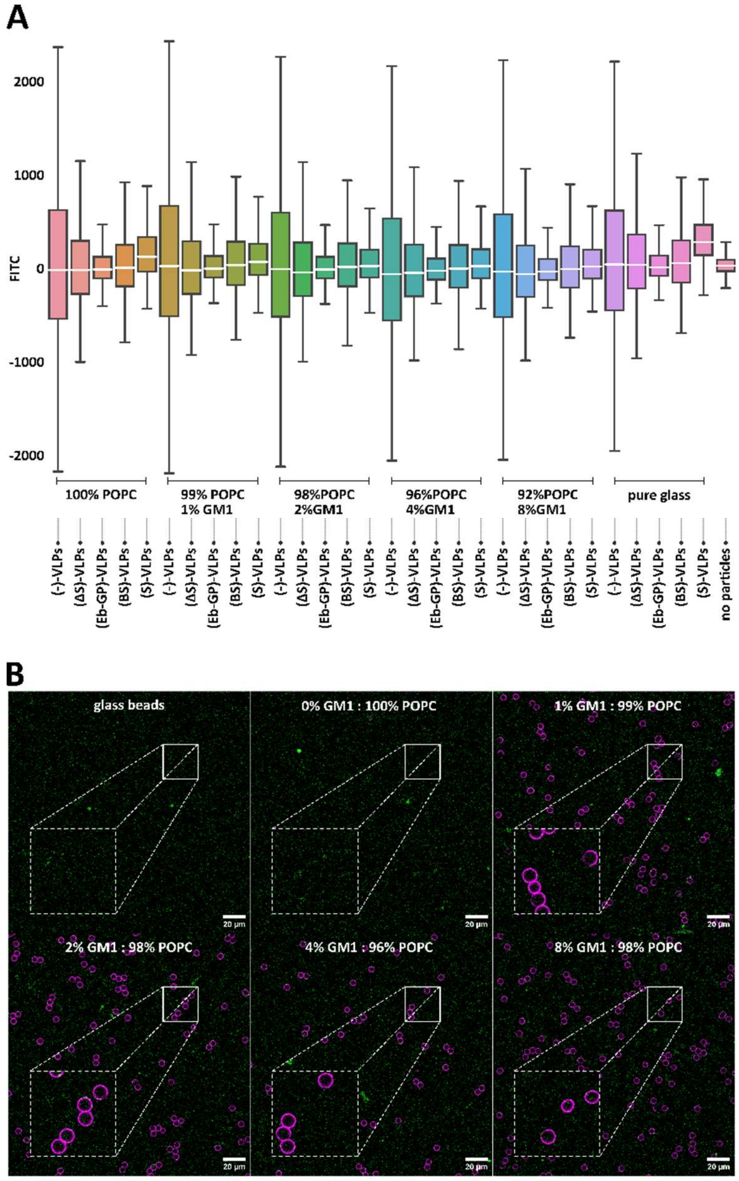


**Fig. S4.**

**GM1-fBSLBs showed no increased binding to VLPs pseudotyped with different viral proteins. A.** Flow cytometry of VLPs pseudotyped with SARS-CoV-2 Spike (S) beta (B), delta (Δ), Ebola glycoprotein (GP) or no protein (-) to BSLBs coated with increasing GM1 concentrations showed no increased binding to GM1. Slightly increased binding was only observed for +S-VLPs to glass beads and POPC coated BSLBs. Box plots showing the quartiles of each data set with N ≥ 10000 with median line in white. **B.** Concentration-dependent insertion of GM1 into fBSLBs was confirmed by cholera toxin B labelling (magenta) and confocal microscopy. Example showing GFP-tagged VLPs (green) pseudotyped with Δ-S.
